# Cryo-EM reveals the structural heterogeneity and conformational flexibility of multidrug efflux pumps MdtB and MdtF

**DOI:** 10.1101/2025.04.25.650598

**Authors:** Surekha Padmanabhan, Clayton Fernando Rencilin, Rupam Biswas, Somnath Dutta

**Affiliations:** Molecular Biophysics Unit, Indian Institute of Science, Bengaluru, India; The Ohio State University, United States

**Keywords:** Efflux Pump, MdtB, MdtF, membrane protein, detergent, *E. coli*, Cryo-EM

## Abstract

RND efflux pumps are a major cause of multidrug resistance in bacteria, particularly in Gram-negative bacteria. They are complex molecular machines forming tripartite assemblies that actively pump out a wide range of antimicrobial agents, including antibiotics, biocides and host defence molecules. However, the presence of multiple RND pumps with overlapping functions in a single bacterium raises questions about their individual functional relevance.

In this study, we determined the Cryo-EM structures of two distinct HAE-RND pumps from *E.coli.,* MdtB and MdtF. MdtF is a unique class of RND pump whose expression is regulated by oxygen availability and is crucial for the survival of *E.coli* in anaerobic growth conditions. Surprisingly, homotrimeric MdtB, previously reported to be inactive, exhibited activity in our experiments. Cryo-EM structure determination of MdtF at 2.8 Å resolution revealed the flexible transmembrane domain with significant conformational changes in the core helixes, leading to the closure of the central pore. While the MdtB structure adopts a resting state, MdtF displays structural dynamics in the presence of DDM. The apo structures of both pumps unveil novel conformational states of HAE-RND efflux transporters. Our findings highlight significant structural and functional divergence among closely related HAE-RND proteins. Despite 71% sequence similarity and phylogenetic proximity to AcrB, the structure of MdtF differs from the solved AcrB structures. In contrast, the MdtB structure exhibits greater resemblance to AcrB, regardless of 32% sequence similarity. Furthermore, our structural analysis provides insights into drug-binding sites and the transport mechanism of these important transporters.

## Introduction

Multiple Resistance-nodulation-cell division (RND) transporters are identified in gram-negative bacteria that can transport a wide range of substrates. The RND family of transporters interacts with the periplasmic membrane fusion protein (MFP) and the outer membrane protein (OMP) to form a macromolecular tripartite complex. The RND pump uses proton motive force as an energy source to efflux substrates (Anes *et al*, 2015). RND pumps are classified based on the type of substrates they transport: hydrophobic and amphiphilic efflux (HAE subfamily) and heavy metal efflux (HME subfamily). *Escherichia coli* (*E.coli*) genome codes for six RND pumps in total; five efflux transporters belong to the HAE-RND family, and one efflux system belongs to the HME-RND family (Anes *et al*, 2015). The HAE-RND efflux transporter of *E.coli* comprises AcrAB, AcrAD, AcrEF, MdtAB, and MdtEF (Tikhonova & Zgurskaya, 2004; Rosenberg, 2000; Baranova & Nikaido, 2002; Nagakubo *et al*, 2002; Kobayashi *et al*, 2006; Nishino & Yamaguchi, 2001). It transports a wide range of substrates like drugs, detergents, dyes, antibiotics, host defence molecules, bile acids and organic. The CusCFBA is the only HME-RND efflux system of *E.coli* that transports Cu and Ag (Moseng *et al*, 2021; Su *et al*, 2011). Among the five HAE-RND pumps, AcrB, AcrD, AcrF and MdtF fall into the Acr cluster (Górecki & McEvoy, 2020). MdtB is the only HAE RND pump from *E.coli* that belongs to the Mdt cluster (Górecki & McEvoy, 2020). It has been reported that the substrate range for all the RND transporters is not very distinct; many RND pumps transport the same substrates. Many of these RND pumps have overlapping functions, while some RND pumps show distinct functions complementing the existing system. AcrB transports a wide range of substrates, while AcrD transports glycoside (Rosenberg, 2000). Furthermore, Growth condition plays a major role in regulating these RND pump expressions and activity (Kobayashi *et al*, 2006). However, very few RND pumps are structurally and functionally characterized.

We performed structural characterization of less explored efflux transporters MdtB from the mdt class and MdtF from the Acr class. Although AcrB and MdtF are from the same class, both the protein activation conditions are different (Zhang *et al*, 2011; Kim *et al*, 2010). They might have an overlapping efflux activity, or their activities can complement in *E. coli*. MdtF transporter is 71% similar to *E. coli* AcrB, whereas MdtB is 32% similar to *E. coli* AcrB. Phylogenetic analysis has revealed that MdtF is closer to AcrB; both are from the Acr class of HAE-RND pumps. Whereas MdtB is from the Mdt class of HAE-RND pumps, it is closer to MdtC (Górecki & McEvoy, 2020).

Among these, the AcrB pump is extensively studied and acts as a benchmark for RND transporters. This pump is ubiquitously expressed and reported to transport multiple substrates. MdtF from RND family show expression and activity in an anaerobic environment. It was found that MdtF is upregulated more than 20-fold in anaerobic conditions (Zhang *et al*, 2011). It actively transports toxic molecules produced in an anaerobic environment to survive extreme conditions (Schaffner *et al*, 2021; Deng *et al*; Zhang *et al*, 2011; Xu & Yan, 2015). Anaerobically active RND efflux transporters are characterized by obligate anaerobes like *Bacteroides fragilis* (Anes *et al*, 2015). Facultative anaerobes like *E. coli* need a specific set of systems to activate and transport substrate in an anaerobic environment apart from the commonly studied aerobic systems. *E.coli,* being a facultative anaerobe and opportunistic pathogen, is a good model organism to study. Additionally, this is found in the human gut and experiences an anaerobic environment. However, there is very little information we have about the RND pumps that coexist and function in both aerobic and anaerobic environments. It may be possible that AcrB and MdtF could show a functional crossover in *E. coli*.

On the other hand, MdtABC is a two-RND subunit system, which has two inner membrane RND transporters, MdtB and MdtC is coded in its operon (Kim *et al*, 2010; Baranova & Nikaido, 2002; Nagakubo *et al*, 2002). MdtB is the only two-RND subunit system in *E. coli*. The homologous two RND transporter systems are identified in *Pseudomonas aeruginosa* (MuxABC), *Salmonella enterica, Serovar Typhimurium, Serratia marcescens, Erwinia amylovora, Pseudomonas putida, and Photorhabdus luminescens* (Górecki & McEvoy, 2020). The two RND transporters, MdtB and MdtC, form an independent homo-trimeric assembly and hetero-trimeric oligomer. Interestingly, the heterotrimeric state MdtB2C is found to be active, whereas homotrimeric MdtB exists in an inactive form (Kim *et al*, 2010). The need for two RND transporters in a system and whether they work together or independently are still unclear. So far, the heterotrimeric RND pump transport mechanism has not been explored much.

The structural architecture of the HAE-RND transporter is conserved. The HAE-RND transporter has three structural domains-docking domain (DD), portor domain (PD) and transmembrane domain (TMD) (Murakami *et al*, 2006). The docking domain is the region of interaction with the membrane fusion proteins (MFP) to form the tripartite complex. All the substrate channels lead to a single funnel/duct in the docking domain region, which connects the outer membrane channel of the tripartite system. Periplasmic domains consist of four subdomains: PN1, PN2, PC1 and PC2 (Murakami *et al*, 2006). These subdomains come together to form highly hydrophobic pockets. The subdomains form two binding pockets in portor domain - access binding pocket (ABP) and deep binding pocket (DBP). PC1 and PC2 form an access binding pocket (ABP) at the periphery of the protein. Followed by, PN1 and PN2 forming a deep binding pocket (DBP) towards the core of the protein. The substrate binding pockets are rich in conserved hydrophobic aromatic amino acids. G loop divides these two pockets and acts as a switch/ valve to close or open the path to DBP from ABP (Eicher *et al*, 2012). The transmembrane domain has 12 transmembrane helixes, and proton gradient-dependent conformational changes are translocated through the transmembrane domain (Zwama & Yamaguchi, 2018).

So far, many conformational states of the RND pump have been captured in its apo form and substrate-bound form. The asymmetric structure of AcrB is solved in the presence of substrates minocycline, doxorubicin, ethidium, dequalinium, detergents (DDM, LMNG, UDM) (Bharatham *et al*, 2021; Tam *et al*, 2021a; Eicher *et al*, 2012; Sennhauser *et al*, 2006; Tam *et al*, 2020; Ababou & Koronakis, 2016) and inhibitors. The asymmetric structure of AcrB reveals that each protomer in the trimer exists in different conformational states, depicting the mechanism of substrate transport. The trimeric HAE-RND transporter undergoes conformational changes in an interdependent cyclic manner to transport the substrate. The protomers are in continuous cooperative movement during substrate translocation, known as a functional rotational mechanism (Tam *et al*, 2021a). The three classical conformational states described by the HAE-RND transporters are access state, binding state and extrusion state in the order of the steps. The movement of the Porter domain is responsible for the distinct conformational difference among these states. Opening and closing of PN1, PN2, PC1 and PC2 are observed in these states (Murakami *et al*, 2006; Seeger *et al*, 2006). In the access conformational state protomer, the protein is ready to take up the substrate from the peripheral opening-access binding pocket and the vestibule formed by PC2 and the transmembrane helix TM 8. It is observed that PC1 and PC2 moved away from each other to open (lose) the access binding pocket. Following that, the protein experiences binding state conformation, in which the substrate enters from the peripheral cavity to the deep binding pocket (DBP), passing the G-loop. Structurally, the subdomains PN1 and PN2 moved away to open the DBP cavity. The central helix from the neighbouring extrusion monomer is inclined to block the exit gap in the DBP of the binding state protomer. Finally, In the extrusion state protomer, the substrate binding pockets and the channel entries are closed. At the same time, the funnel duct is opened because of the movement of the central helix to transport the substrate. The access binding pocket and the deep binding pockets are closed by the movement of PC1 & PC2 and PN1 & PN2. The TM8 is moved closer to PC2 in extrusion conformation to close the vestibule entry (Murakami *et al*, 2006; Seeger *et al*, 2006).

## Results

In this study, we aim to characterize the structure and function of two efflux pumps, MdtB and MdtF, from *Escherichia coli*, which are involved in the extrusion of antibiotics. These two HAE-RND family transporters have been poorly characterized to date, and no structural information is currently available. In addition to elucidating their individual structures, we are interested in understanding how these pumps may interact or coordinate in the process of drug efflux. Preliminary computational studies and biochemical characterization of the MdtB and MdtF efflux pumps are carried out to gain insights into their overall architecture and guide subsequent experimental approaches.

### Bioinformatic and Biochemical Characterization of MdtB and MdtF

To begin our analysis, we performed sequence alignments of efflux pumps MdtF and MdtB with transporter AcrB. Pump MdtF shares 71% sequence identity with AcrB, whereas MdtB shares only 32% identity with AcrB, indicating that MdtF and AcrB are highly conserved, while MdtB is divergent (**Figure 1A**). We specifically analyzed conserved amino acid residues across the three transporters. Notably, Asp431, Asp432, and Lys938 (MdtF)-as well as Phe611-are conserved in all three proteins (**Supplemental Figure 1**). Among these, the three charged residues (Asp431, Asp432, and Lys938) are known to be essential for proton translocation, suggesting functional conservation in the mechanism of transport.

**Figure 1:**
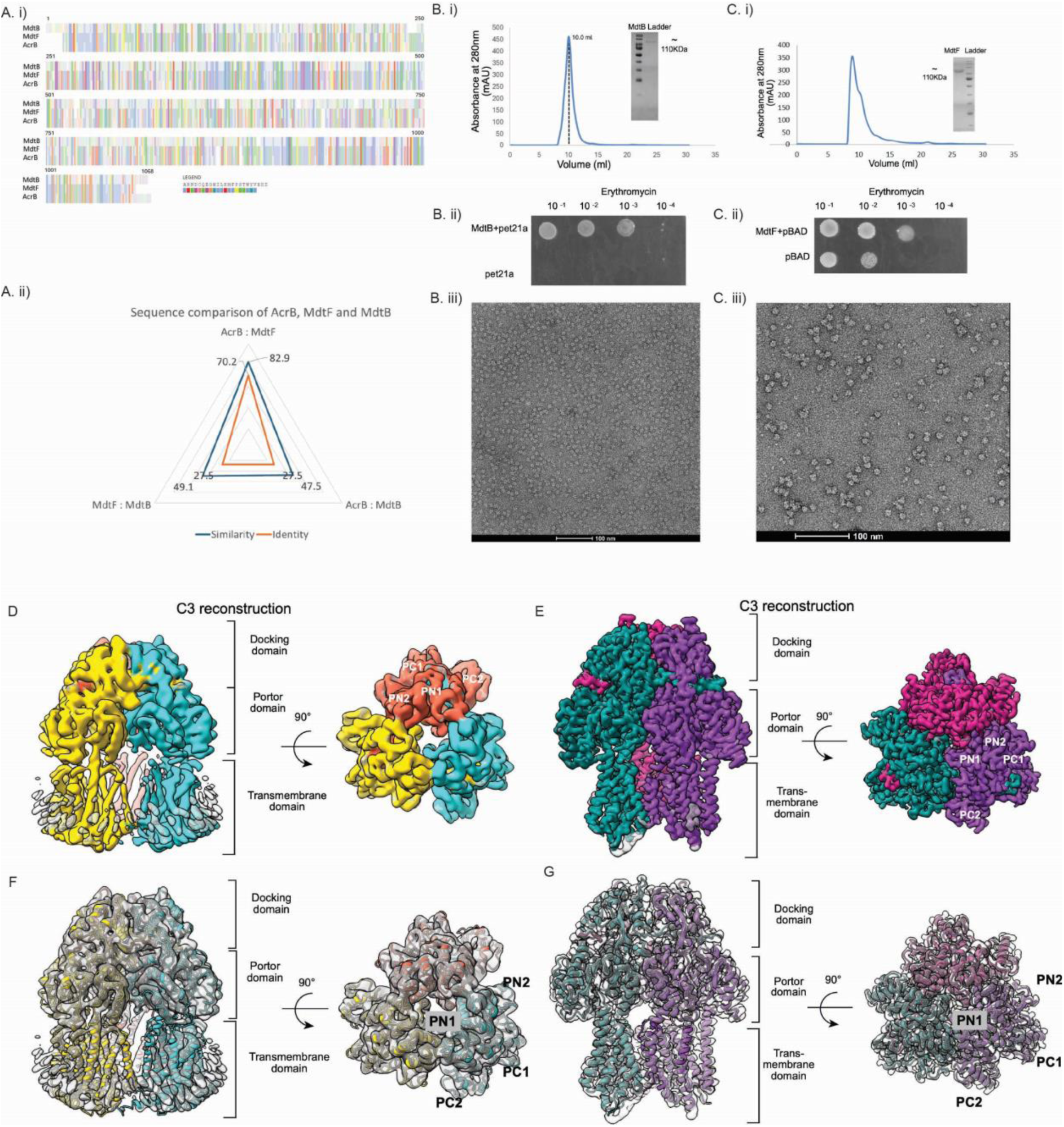
Biochemical and structural characterisation of MdtB and MdtF. **A. i)** Multiple sequence alignment of RND pumps MdtB, MdtF and AcrB. **A. ii)** Sequence comparison analysis displaying similarity and identity between the RND efflux pumps MdtB and MdtF with the reference pump AcrB. **B. i)** SEC elution of MdtB in the S200 column at 10 ml volume corresponding to the trimeric molecular weight of the protein. Eluted MdtB sample ran in the SDS PAGE gel (inside) **C. i)** SEC elution of MdtF in the S200 column at 10 ml elution volume corresponding to the trimeric molecular weight and its SDS PAGE of purified MdtF (inside). **B. ii), C. ii)** Agar plate assay showing the in vivo activity of MdtB and MdtF, respectively, in the presence of erythromycin substrate. **B. iii), C. iii)** Negative staining TEM (NSTEM) micrograph of MdtB and MdtF sample. **D.** Side view and top view of symmetric MdtB Cryo-EM density map resolved with C3 symmetry at a global resolution of 4.3 Å. Each protomer is coloured differently in the Cryo-EM density **E.** Fitting of the built atomic model of MdtB in the Cryo-EM density map. **F.** side view and top view of MdtF Cryo-EM map resolved at a 2.8 Å resolution in C3 symmetry. Each protomer is coloured differently in the Cryo-EM density. **G.** Fitting of the built atomic model of MdtF in the Cryo-EM map.

To validate the functional activity of efflux pumps MdtB and MdtF prior to structural characterization, we performed *in vivo* agar plate-based growth assays for both transporters. As expected, *E.coli* cells harbouring the MdtF-expressing vector exhibited active efflux of erythromycin and survived in the presence of the antibiotic. Notably, survival was observed even at erythromycin concentrations of ∼4 µM, indicating that MdtF is functionally active (**Figure 1C ii**). Interestingly, when the same assay was performed with pump MdtB, the homo-trimeric form of MdtB also demonstrated the ability to transport erythromycin (**Figure 1B ii)**. This observation is particularly significant, as previous studies have suggested that the hetero-trimeric assembly of MdtB_2_C forms an active efflux channel, while the homo-trimeric MdtB alone was thought to be inactive. Our findings, therefore, provide the first evidence that the homo-trimeric MdtB can function as an active channel under these conditions. To further confirm these results, we conducted a liquid media growth assay using both MdtB and MdtF. Consistent with the agar plate assay, both pumps exhibited efflux activity against erythromycin, supporting their functional roles in antibiotic resistance.

To facilitate structural studies, efflux pumps MdtF and MdtB were purified following detergent extraction with 2% n-dodecyl-β-D-maltoside (DDM), with the concentration subsequently reduced to 0.03% for downstream purification. A two-step purification strategy was employed, beginning with immobilized metal affinity chromatography (IMAC), followed by size-exclusion chromatography (SEC). Both proteins eluted at approximately 9 mL on a Superdex™ 200 Increase column (**Figure 1B.i & 1C.i**), consistent with the trimeric form. SDS-PAGE analysis confirmed the purity and homogeneity of the samples, with a distinct band observed at ∼110 kDa (**Figure B.i & C.i**). To further examine their oligomeric state, negative-stain transmission electron microscopy (TEM) was performed. The TEM micrographs revealed that both MdtF and MdtB assemble as homotrimers (**Figure 1B iii & 1C iii**), in agreement with the SEC elution profiles. These results, together with the high expression yields and stable trimeric assemblies, provided a strong foundation for advancing to cryo-electron microscopy (cryo-EM) studies.

### Structural characterization of MdtB using single particle Cryo-EM

Negative-stain transmission electron microscopy (TEM) was initially employed to evaluate the structural integrity and cryo-EM suitability of the purified MdtF and MdtB proteins. The collected micrographs confirmed that both samples were uniform and monodispersed, ideal for cryo-EM structural study (**Figure 1B iii & 1C iii**). Therefore, we proceeded to collect extensive cryo-EM datasets for both transporters to enable structural determination. The collected cryo-EM data were processed to generate reference-free two-dimensional (2D) class averages, which displayed characteristic top and side views consistent with HAE-RND family transporters (**Supplemental Figure 2**). Further, we performed a three-dimensional (3D) reconstruction of transporter MdtB, yielding a map at an overall resolution of ∼4.3 Å (**Figure 1D**). This resolution was sufficient to confidently trace the polypeptide backbone and construct a reliable atomic model of MdtB (**Figure 1E**).

We successfully resolved the high-resolution cryo-EM structure of MdtB, revealing an overall fold characteristic of the HAE subfamily within the RND transporter family. Notably, this represents the first high-resolution structural determination of MdtB. The reconstruction allowed us to clearly visualize all transmembrane helices within each protomer, as well as the complete periplasmic domain. The homotrimeric assembly of MdtB closely resembles that of canonical RND transporters, such as AcrB from *E. coli* (**Figure 1D&F**). Structurally, each protomer of MdtB comprises three major domains: the docking domain, the porter domain, and the transmembrane (TM) domain (**Figure 1D&F**). The docking domain, located in the periplasm, contains two subdomains-DN1 and DN2-that form a central funnel-like structure (**Figure 2A)**. The DN1 and DN2 subdomains of the docking domain form a central funnel that gathers substrates from multiple entry pathways and channels them outward via the tripartite efflux complex. The Porter domain of MdtB is situated in the periplasmic region, in between the docking and transmembrane (TM) domain and consists of four subdomains: PC1, PC2, PN1, and PN2 (**Figure 2A**). The fold of each domain is identical to the RND subfamily of proteins, and each has β-ɑ-β sandwich motif (**Figure 2D**). These domains come together to form the substrate-binding pockets, which allow these RND pumps to efflux the antibiotics and drugs for recognizing and extruding a wide range of antibiotics and toxic compounds.

**Figure 2:**
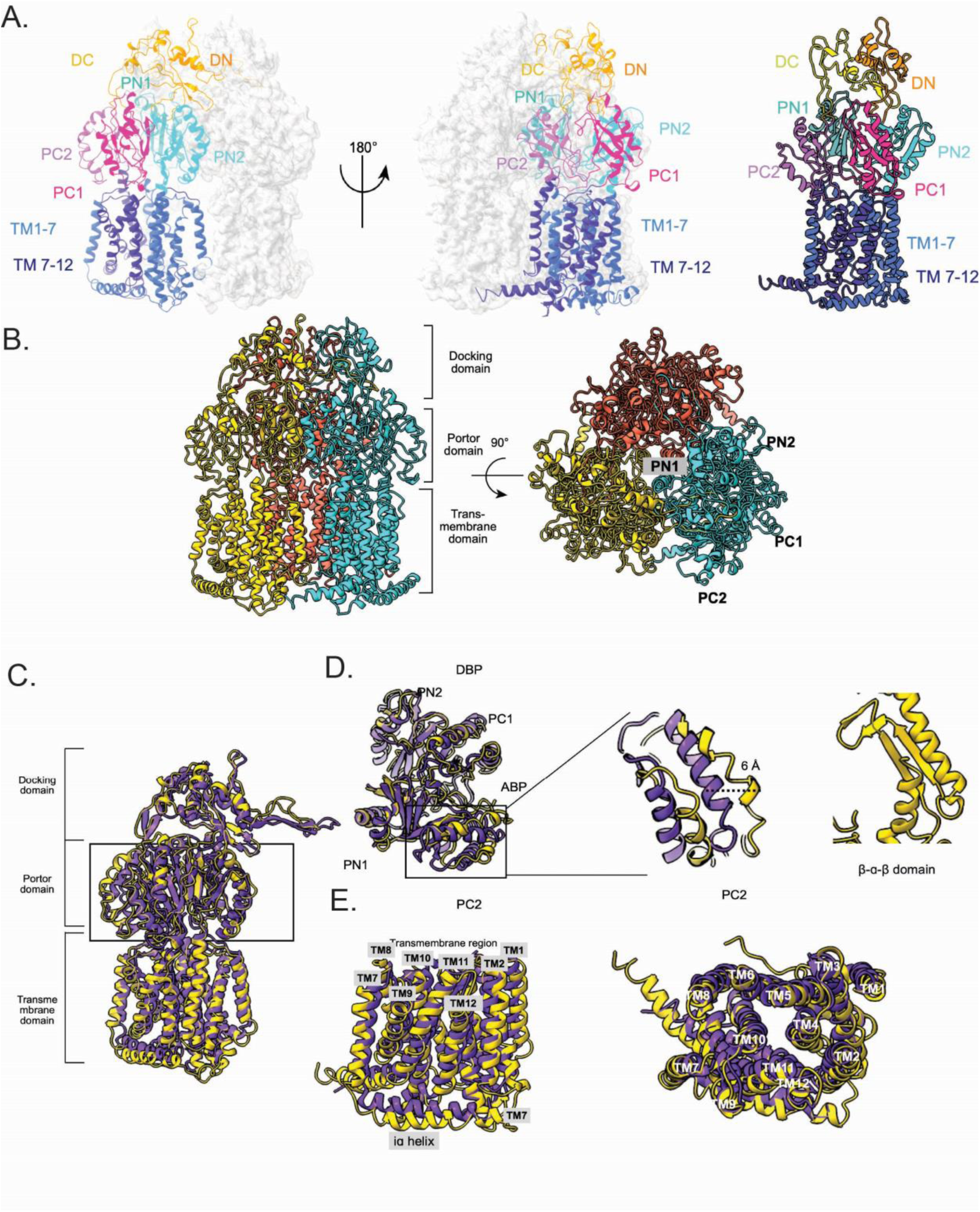
Atomic structure of MdtB and structural comparison with AcrB. **A.** Monomeric model of MdtB monomer built from the Cryo-EM density map. Sub-domains are highlighted in varied colours and labelled. **B.** The atomic model of the MdtB homo-trimer shows its tertiary architecture from the side and top. **C.** MdtB and AcrB protomers are superimposed; yellow represents MdtB, while purple shows AcrB. **D.** Comparison of the Porter domain. Movement in the PC2 domain is highlighted. **E.** Transmembrane domain helixes are compared between MdtB and AcrB.

To assess conformational variability within individual protomers of MdtB, we conducted an asymmetric reconstruction of protein MdtB. The resulting map showed that the overall trimeric architecture remained consistent. Superimposition of the three protomers from the asymmetric reconstruction demonstrated a high degree of similarity, indicating minimal structural heterogeneity between the protomers. For further comparison, individual protomers of MdtB were aligned with the three known conformational states of AcrB protomers **(Figure 2D).** Notably, this analysis revealed that the Access Binding Pocket (ABP) in MdtB adopts a closed conformation. Specifically, the PC2 subdomain was found to shift approximately 6 Å toward the PC1 subdomain, thereby narrowing the ABP (**Figure 2D & 5A**). The conformational states of the PN1 and PN2 subdomains in MdtB were consistent with the binding state of RND transporters. However, the PN2 subdomain was displaced outward relative to PN1, resulting in the formation of a wider deep binding pocket (DBP) (**Figure 3**). Structural comparison with the binding state of AcrB revealed a high degree of similarity, with a main-chain root mean square deviation (RMSD) of 1.421 Å upon superposition (**Supplemental figure 8**). Additionally, the transmembrane domain of MdtB displayed a characteristic pseudo-two-fold symmetry, with TM helices 1–7 and 8–12 forming two structural repeats within the membrane-spanning region.

**Figure 3:**
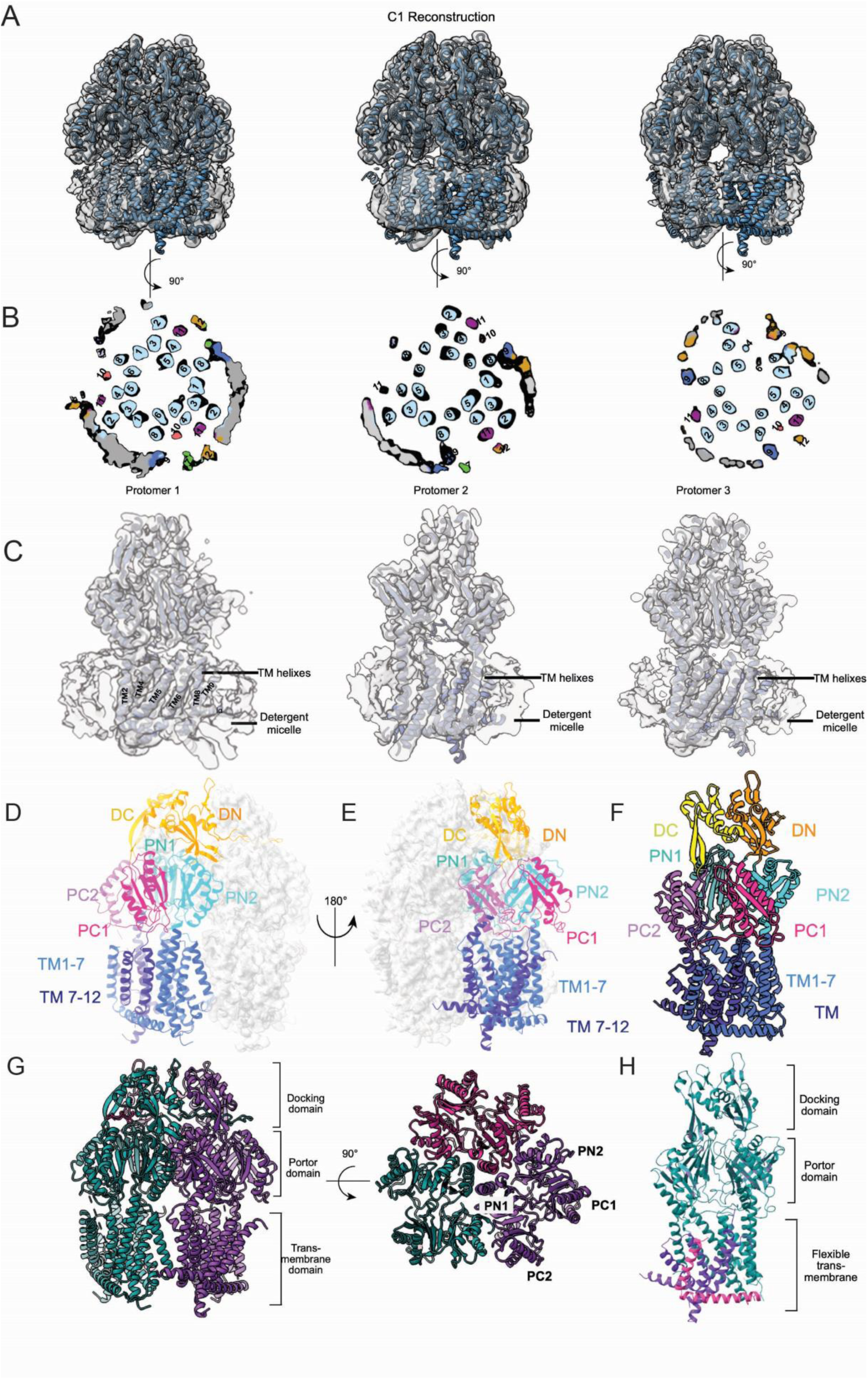
Asymmetric Cryo-EM density maps display peripheral helix flexibility. **A.** 3D classes of MdtF fitted with MdtF model. **B.** The cross-sections of the top view are shown for a better view of all the transmembrane helices. In class 1 protomer1, all 12 transmembrane helixes are resolved, while in protomers 2 and 3, some TM helixes are missing. Peripheral helixes showing flexibility are highlighted in different colours. The presence and absence of helixes are highlighted in each protomer of the three classes. **C.** Side view of atomic model fitted onto cryo-EM map, showing the presence or absence of peripheral helices. **D, E, F.** The monomeric model of MdtF was built from the asymmetric cryoEM map. Each domain is highlighted in a variant colour. **G.** Side view and Top view of MdtF trimeric model. **H.** Comparison of all the three protomers by superimposing. The flexible transmembrane helixes are highlighted in purple and magenta colour.

The overall conformations of the transmembrane helices and the arrangement of the core helices were largely consistent with those observed in the prototype AcrB structure, although notable positional shifts were detected. These included significant displacements in the iα helix, the loop connecting to the horizontal helix, specific segments of TM12, and the TM1 helix in MdtB (**Figure 2C**). Notably, TM helix 8 adopted a folded conformation at the interface between the transmembrane and periplasmic domains, showing a displacement of approximately 5 Å relative to the access-state protomer of AcrB. Furthermore, TM helices 12 and 1, along with the horizontal helix, displayed shifts in the range of 2–2.5 Å when compared to the corresponding regions in AcrB (**Figure 2E**).

### Cryo-EM analysis of MdtF shows the flexibility of the transmembrane domain

As outlined earlier, the trimeric structure of MdtF was initially evaluated using negative-stain transmission electron microscopy (TEM) (**Figure 1C.(iii)**). To further characterize its structure at high resolution, we conducted cryo-electron microscopy (cryo-EM) analysis. A total of 4,000 movie frames were acquired, and three-dimensional reconstruction applying C3 symmetry produced a map of MdtF at an overall resolution of approximately 2.8 Å, as determined by the gold-standard Fourier shell correlation (GS-FSC) at the 0.143 threshold (**Supplemental Figure 3**). This resolution permitted reliable model building, which was performed using *Phenix* and *Coot*. The resulting atomic model of MdtF revealed an overall fold characteristic of the HAE subfamily of RND transporters, including AcrB, MexB, AdeB, BpeB, and AcrD (**Figure 3 and Figure 3G**). The C3-symmetric density map resolved the core transmembrane (TM) helices in all three protomers (**Figure 1G**). Notably, peripheral TM helices (TM7 and TM9–12), which interact with the surrounding DDM micelle, remained unresolved, likely due to flexibility.

Interestingly, asymmetric reconstruction of MdtF revealed density corresponding to the peripheral transmembrane helices, which were not observed in the C3-symmetrized map **(Figure 3B).** In one of the three protomers, all 12 transmembrane helices were clearly resolved. However, the remaining two protomers still the full TM helix set is not resolved, even after applying asymmetric processing (**Figure 3B**). Although peripheral transmembrane helices could be visualized, their densities appeared at a noticeably lower resolution than those of the central core helices (**Supplemental Figure 3**). These observations indicate that the three MdtF exhibit different conformational behaviours, and not all 12 transmembrane helices are resolved simultaneously in every protomer. The central helices (TM1–6, TM8, TM10 and TM12) were consistently well resolved across all 3D classes analyzed (**Figure 3B**). In contrast, the peripheral helixes TM7, TM9, and TM11 resolved variably depending on the protomer and classification state. Notably, while some peripheral helices such as TM7, TM9, and TM11 could be resolved individually, they were never visualized concurrently in a single protomer. For instance, in class 1, protomer 1 displayed electron density for both TM7 and TM12, whereas these helices were absent in the other two protomers (**Figure 3B**). In class 3, only TM9 was resolved in one protomer, while the remaining helices were not visible. Remarkably, in class 2, protomer 1 exhibited density for all three peripheral helices (TM7, TM9, and TM11)simultaneously (**Figure 3B**). The variability observed in resolving the three peripheral transmembrane helices across different protomers highlights the inherent flexibility within the transmembrane domain, which likely contributes to the difficulty in capturing all helices simultaneously in a single reconstruction (**Figure 3B**). Additionally, local resolution analysis of the final cryo-EM map revealed that the peripheral helices were resolved at a lower resolution of approximately 5 Å (**Supplemental Figure 3**), suggesting increased flexibility compared to the more stable core helices. Moreover, the traced transmembrane helices exhibited substantial conformational deviations compared to other characterized RND transporters.

### Structure comparison between AcrB and MdtF

To analyse structural differences among RND transporters, we conducted a comparative analysis of MdtF with both AcrB and MdtB, revealing distinct variations across all major subdomains. Initially, we examined the conformational states of the Porter domain by aligning the periplasmic region of the MdtF protomer with the three known conformational states of the *E. coli* AcrB protomer (**Supplemental Figure 8**). This superimposition revealed notable structural differences, which are discussed in the following sections.

AcrB is known to cycle through three distinct conformational states: the access, binding and extrusion states. In our study, cryo-EM analysis yielded a single high-resolution structure of MdtF. To explore its conformational state, we compared the periplasmic domain of MdtF structure with the three protomer states of AcrB. The structural alignment and root mean square deviation (RMSD) calculations were performed to identify the similarity between the periplasmic domains. The RMSD between MdtF and the binding-state protomer of AcrB was approximately 1.08 Å, indicating the closest structural resemblance. In contrast, comparisons with the access and extrusion states yielded RMSD values of ∼1.156 Å and ∼1.133 Å, respectively (**Supplemental Figure 8**). These results suggest that the resolved MdtF structure most closely resembles the binding conformation of AcrB. Notably, domain-wise superposition of PC1, PC2, and PN1 yielded RMSD values around ∼1 Å, demonstrating a high degree of structural conservation in these subdomains (**Figure 4A**). Structural analysis further revealed that the PN2 domain helix in MdtF is shifted approximately 3.7–4 Å closer to the PN1 domain when compared to the binding-state protomer of AcrB (**Figure 4B**). This inward movement of PN2 appears to close the channel, resulting in a reduced deep binding pocket (DBP) volume in MdtF, though it is wider than that observed in AcrB’s access state. Notably, all three MdtF protomers adopt a similar conformation, ensuring that the orientation of the central helices does not block the DBP exit of a neighbouring protomer. These observations indicate that the Porter domain of MdtF likely represents an intermediate state transitioning from the canonical binding conformation.

**Figure 4:**
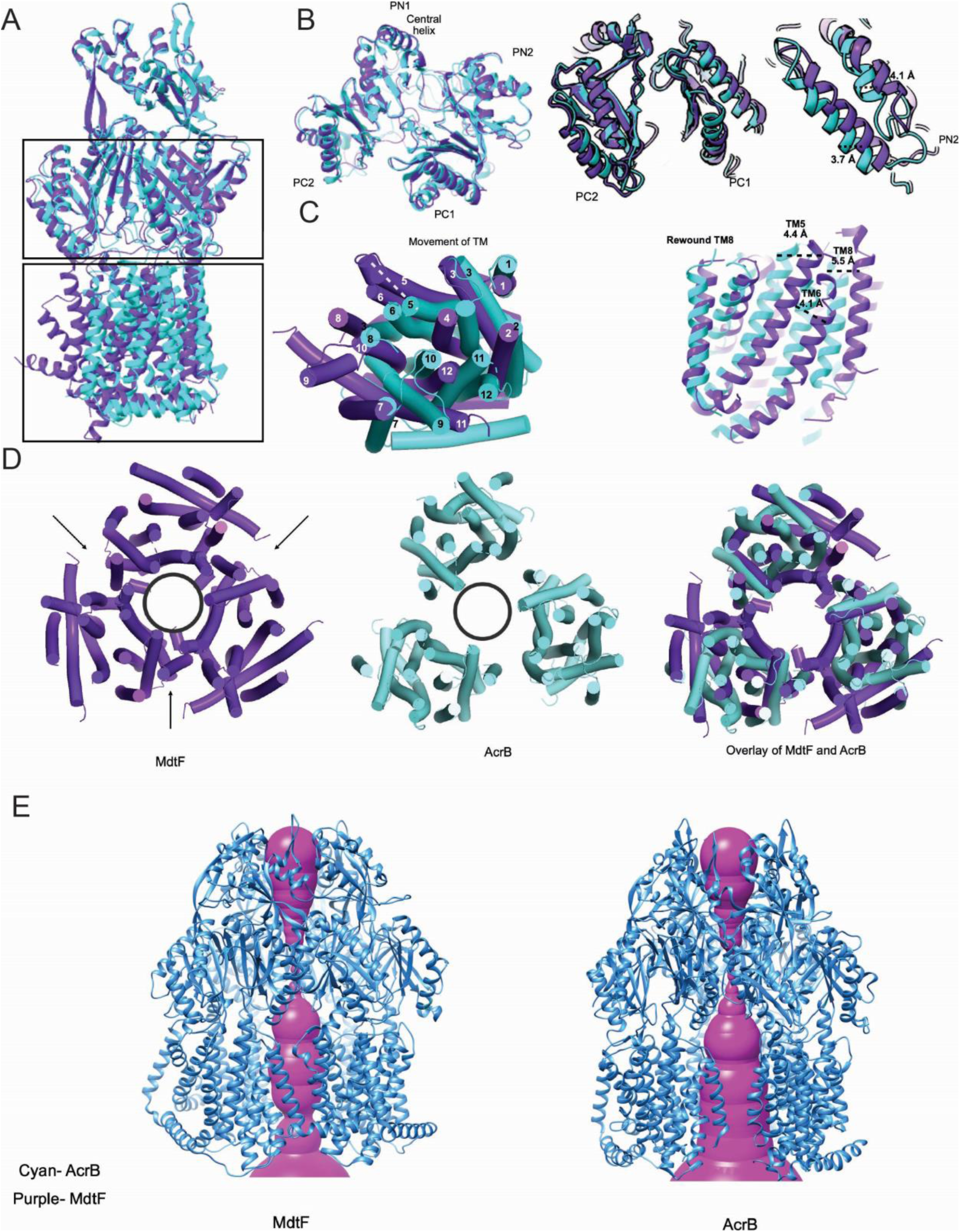
Structural comparison of MdtF with AcrB. **A.** MdtF and AcrB monomers are superimposed. **B.** The periplasmic region is aligned with the binding state protomer of AcrB. Shift in the PN2 domain helix is observed. **C.** Movement in the transmembrane helixes are highlighted. Transmembrane helix 5, 6 and 8 are the core helix that shifts towards the pore. **D.** The Central pore of MdtF and AcrB was compared. movement of core helixes in MdtF caused the central pore to close from the transmembrane region. **E.** Pore analysis is performed with MOLEonline web tool. Shrinkage of the pore is observed in the transmembrane region.

The high-resolution structure of MdtF offers valuable insights into its three-dimensional architecture and highlights key structural differences from AcrB, particularly within the transmembrane (TM) region. The most prominent variations were observed in the core helices of the TM domain. Consistent with prior reports describing a pseudo-two-fold symmetry in the TM subdomains of RND transporters, our analysis revealed that MdtF also exhibits this architectural feature. Specifically, helices TM1–7 and TM8–12 form two symmetrical halves of the transmembrane domain. MdtF contains a total of 12 TM helixes, TM 1-6 and TM8 are assembled as the core helixes of the RND pump, while TM7 and TM9–12 form the peripheral helix (**Figure 3H**).

Structural superimposition of MdtF with the binding-state conformation of AcrB revealed substantial displacements in several transmembrane helices. A detailed comparison of the TM subdomain showed that helices TM4, TM5, TM6, and TM8 in MdtF were shifted approximately 6–7 Å toward the central pore (**Figure 4C**). The four helixes were squeezed towards the centre while the remaining helixes were twisted and moved away to facilitate the core helix movement. TM1, TM2, and TM3 of MdtF moved away from the center compared to AcrB. In previously characterized RND pumps, TM8 undergoes conformational changes, unwinding in the access state and rewinding in the extrusion state, to regulate vestibule opening for substrate entry. In MdtF, however, TM8 remains fully folded in an α-helical conformation and is angled in a manner that creates a widened vestibular space (**Figure 5B**).

**Figure 5:**
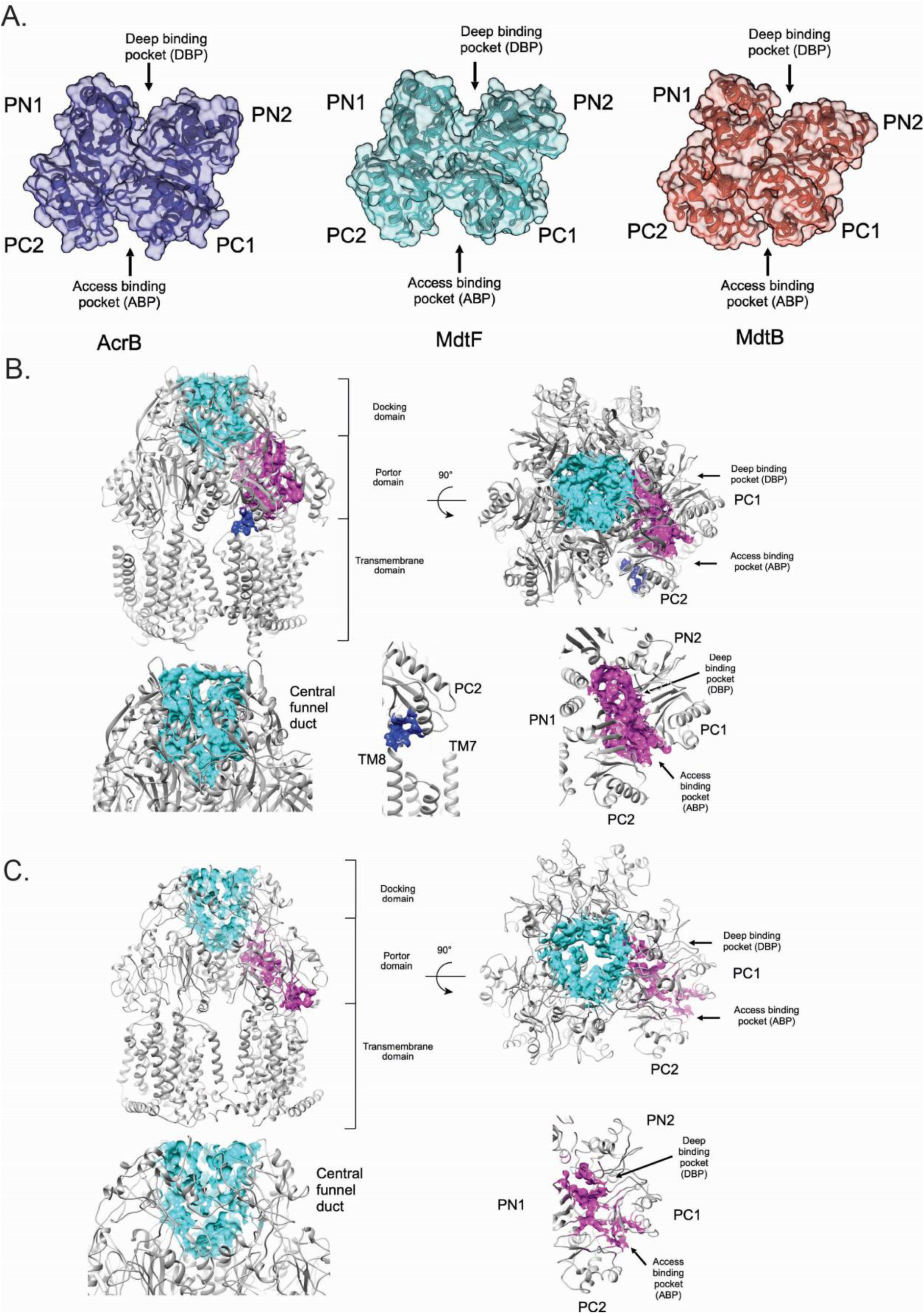
Comparison of binding pockets and substrate transport channels of MdtF and MdtB. A. Access and deep binding pockets of AcrB, MdtF and MdtB are highlighted and sizes are compared. MdtF access binding pocket is wider compared to other proteins. MdtB binding pockets are narrower than AcrB and MdtB. B. Substrate transport channels of MdtF are highlighted in different colours. Access binding and deep binding pocket channels are wide open, and TM8 is in the rewound state with the vestibule opening. **C.** Substrate transport channels of MdtB are highlighted in different colours. Narrower access and deep binding pockets are highlighted.

Furthermore, the cryo-EM map of MdtF revealed a well-defined density corresponding to DDM molecules within the transmembrane domain. These molecules are located in specific binding pockets, notably in the groove between TM8 and TM6 and within the crevice formed by TM1 and TM2 (**Figure 6**). The modelled structure indicates that hydrogen bonding with polar residues in these regions stabilizes the DDM molecules, suggesting a potential role for detergent interactions in maintaining the structural integrity of the transmembrane helices.

**Figure 6:**
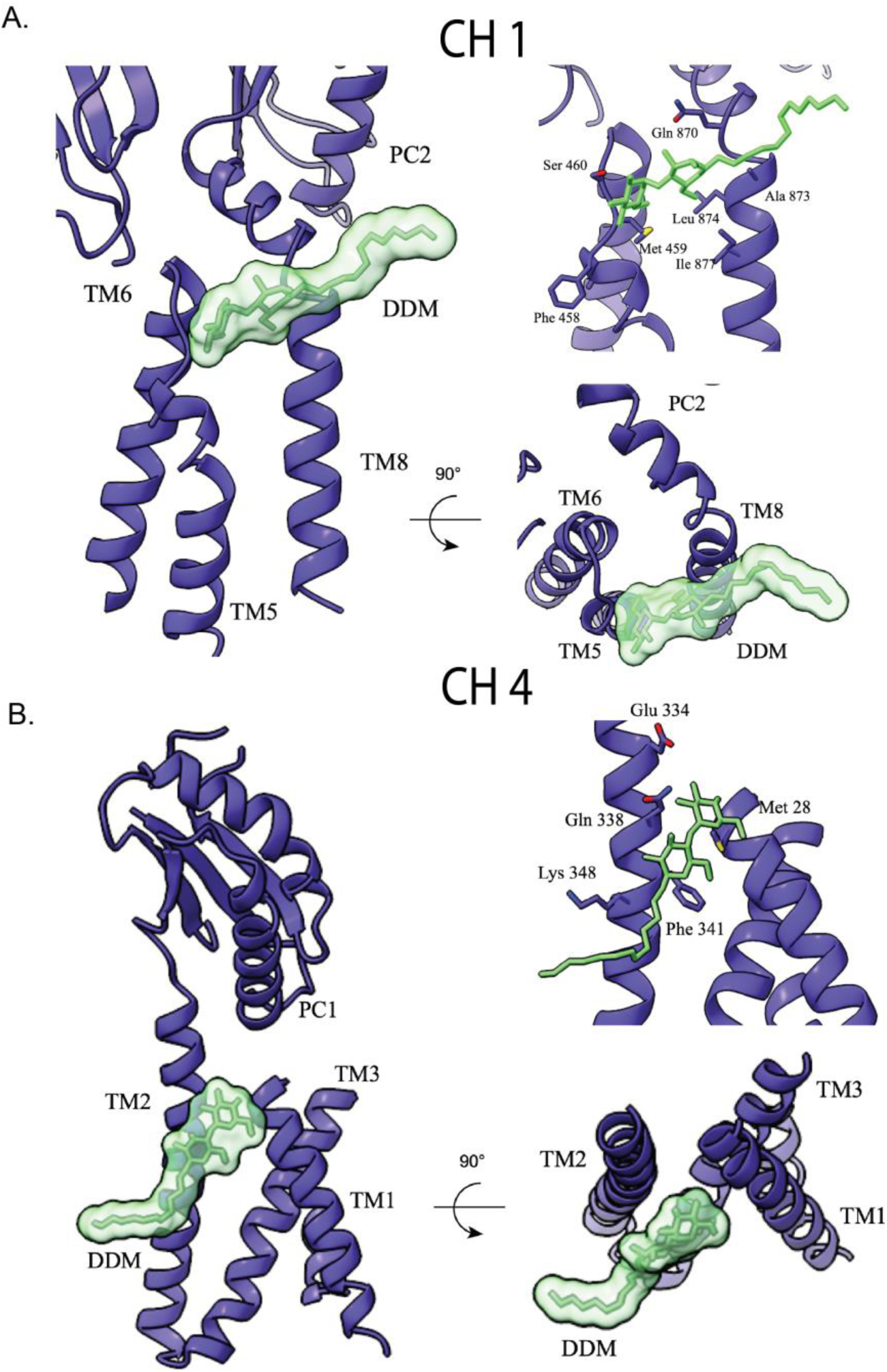
DDM bond in CH1 and CH4 channel groove. A. DDM binding to near the TM8 vestibule entry. B. DDM binding in the TM1/TM2 groove of the MdtF.

### Structural differences between the substrate binding pockets of AcrB, MdtF and MdtB

Structural analysis of MdtB and MdtF revealed that both adopt conformations distinct from the three classical states typically described for RND transporters. In RND pumps, substrates initially access the Porter domain through the access binding pocket (ABP), which is formed by the PC1 and PC2 subdomains. Followed by, substrates are translocated toward the deep binding pocket (DBP), crossing the gate loop (G loop), which functions as a selective barrier. This switch or gating loop acts as a valve, controlling the entry of substrates into the DBP. The G loop is inherently flexible due to the presence of glycine residues, and this flexibility is critical for the dynamic opening and closing of the entry channel, thereby regulating substrate passage into the DBP.

To further explore the structural basis of substrate gating, we examined the gating loop (G loop) in both MdtF and MdtB. The G loop in MdtF spans residues 611–621 (GGFGFSGQGQNN) and contains five glycine residues, one more than the corresponding loop in AcrB, suggesting enhanced flexibility (**Supplemental Figure 7B**). This increased flexibility likely facilitates the loop’s dynamic movement during substrate translocation. In the access state of RND transporters, the G loop faces inward, blocking the deep binding pocket (DBP), whereas in the binding and extrusion states, it reorients away from the DBP, enabling substrate entry. Consistent with this, the G loop in MdtF adopts a conformation that resembles the binding state, thereby exposing the DBP. In contrast, the G loop in MdtB comprises residues 609–619 (VGVDGTNPSLN) and has a reduced glycine content, resulting in a more rigid conformation. In our structure, this loop appears displaced upward, further supporting its limited flexibility (**Supplemental Figure 7B**). Importantly, phenylalanine at position 615 is conserved across RND pumps, including MdtF, but is absent in MdtB(**Supplemental Figure 1**). Furthermore, both binding pockets are surrounded by highly conserved aromatic residues, which are likely important for substrate recognition and stabilization.

Furthermore, we examined the structural and morphological features of the substrate transport channels in MdtB and MdtF using the MOLE and CASTp web tools (Tian et al., 2018; Berka, 2012). The regulation of substrate entry at the vestibule is governed by the dynamic movements of the PC2 domain and the conformational change of TM8, which undergoes folding and unfolding transitions (**Figure 5B & Supplemental Figure 7D**). These conformational changes are closely linked to the functional rotational mechanism characteristic of substrate transport in RND pumps. In the access state, TM8 unwinds at the interface between the periplasmic and transmembrane domains and shifts away from PC2, thereby opening the entry pathway. In contrast, during the extrusion state, TM8 refolds into an α-helix, sealing the gap between PC2 and TM8 and effectively closing the channel (**Figure 5B & Supplemental Figure 7D**).

In the case of MdtB, the TM8 helix appears re-folded, effectively sealing the vestibule entry like the extrusion state. Additionally, the PC2 domain is shifted toward PC1 (**Figure 5A & Supplemental Figure 7D**). This configuration closely reflects the extrusion state; however, TM8 and PC2 in MdtB are positioned approximately 3 Å closer together compared to their counterparts in the extrusion state of AcrB (**Supplemental Figure 7D**). In contrast, although the PC2 domain of MdtF adopts an open conformation, the TM8 helix remains fully folded, thereby occluding the vestibule entry while leaving the access binding pocket (ABP) entry unobstructed (**Supplemental Figure 7D**). To better understand the functional implications of the observed conformational state, we analysed the MdtF channel and found that the central pore is closed at the transmembrane region (**Figure 4E**). Specifically, the core transmembrane helices (TM1 to TM6 and TM8) from each protomer are inclined inward and move towards the centre. The inward displacement of transmembrane helices creates a narrowed passage that appears to seal the central channel (**Figure 4E**). As a result, the radius of the bottom opening in MdtF is reduced by approximately 6 Å compared to that of AcrB. These observations suggest that while MdtF periplasmic domain adopts a conformation closely resembling the substrate-binding state of AcrB, its transmembrane closure likely hinders substrate transportation under this conformation. Extending this observation, structural superimposition of MdtB and MdtF offered additional insights into their distinct substrate binding pocket configurations. The access binding pocket (ABP) in MdtB was narrower than in MdtF, due to a ∼6 Å shift of the PC2 domain towards PC1 (**Supplemental Figure 7C**). In contrast, the deep binding pocket (DBP) of MdtB was wider than that of MdtF, with the PN2 domain shifted away from PN1. In the apo form of MdtB, both the vestibule and ABP were closed, while the DBP remained open. On the other hand, MdtF displayed a wider ABP due to the outward displacement of the PC1 and PC2 domains. The ABP in MdtF was broader than in the access-state conformation of other RND pumps and closely resembled the binding-state configuration (**Figure 4B & 7A**).

## Discussion

In this current study, we employed biochemical and cryo-EM-based structural analyses to characterize the efflux pumps MdtB and MdtF under the superfamily of HAE-RND (resistance-nodulation-division) pumps from *Escherichia coli.* Multiple sequence alignment of HAE-RND pumps displayed a low percentage of sequence similarity between MdtB and AcrB (32%), while a high percentage of sequence similarity is observed between MdtF and AcrB at 71% (**Figure 1A & Supplemental Figure 1**). Moreover, MdtB and MdtF are farther in the phylogenetic analysis and belong to two different clusters, Mdt and Acr, respectively (Górecki & McEvoy, 2020). Among the five HAE-RND pumps, AcrB, AcrD, AcrF and MdtF fall into the Acr cluster (Górecki & McEvoy, 2020). MdtB is the only HAE RND pump from *E.coli.* that belongs to the Mdt cluster (Górecki & McEvoy, 2020).

We performed structural characterization of less-explored efflux transporters MdtB from the mdt class and MdtF from the Acr class. Our findings reveal that these RND pumps are captured in structurally distinct states. Although AcrB and MdtF are from the same class, both the protein activation conditions are different (Zhang *et al*, 2011; Kim *et al*, 2010). They might have an overlapping efflux activity, or their activities can complement in *E. coli*. Previous studies show that MdtB is active in heterotrimeric arrangements. Our functional data demonstrate that homotrimeric MdtB could also be an active RND pump in *E. coli*.

The solved MdtF structure is a symmetric trimer; upon performing a structural comparison with other RND pumps, we observed that the MdtF structural state is close to the binding state of AcrB (PDB ID: 2dr6). The periplasmic domain (PD) structural features resemble the binding state (T state) with open ABP, wider DBP and G-loop facing outside. Also, the conformation of the G-loop resembles the binding state protomer, which is outward-facing.

Interestingly, the TMD structural state is very different from that of the reported RND pumps. In the cryo-EM map of MdtF, it was also observed that the three peripheral helixes are solved at low resolution in an asymmetric protomer and couldn’t capture all the peripheral helixes (TM7, TM 9, TM 11) when symmetry is applied (**Figure 3**). Asymmetric reconstruction demonstrates the flexibility of TM domains of individual protomers within MdtF, where all 12 transmembrane helixes are captured in protomer-1 and nine TM helices in protomer-2 and −3 (disordered TM7, TM9 and TM11) of MdtF (**Figure 3**). These observations suggested that the peripheral helixes were flexible, which could be challenging to capture in the structure. Since the protein is in its native environment and is in constant movement before freezing, it is not in a static state. Therefore, heterogeneity in the sample is best captured using the cryo-EM technique. The discreet heterogeneity in the sample can be captured by performing 3D classification, which clusters the discreet groups of samples from a dataset in different classes. Whereas, if the heterogeneity in the sample is continuous, it is hard to capture by clustering into groups. The structure comparison shows that core TM helices TM 4, 5, 6, and 8 are moved towards the central pore, causing the pore to close. The TM helixes of MdtF show a major global shift in the position of our structure, the TM helixes are displaced ∼4.1Å and 5.5Å toward the centre. Many such transient conformations of RND pumps have been solved using the cryo-EM technique. In the Adeb structure, intermediate states of the transport mechanism were solved using a cryo-EM technique, where the opening of the transmembrane domain is observed (Morgan *et al*, 2021).

In RND pumps, four channels are identified to transport different types of substrates (CH1-CH4) (Zwama & Yamaguchi, 2018; Tam *et al*, 2021b). Among the four channels, three of the channel entries are from the transmembrane domain. The channel 1 (CH1) entrance is located at the vestibule between TM8 and PC2, opening to the cell membrane. Meanwhile, the CH4 entrance is between the TM1/TM2 cleft. Both have common substrates, such as β-lactam, fusidic acid, and DDM. The CH3 entrance faces the central pore, which is specific to small polar aromatic molecules, and the CH2 opening is via ABP. Although substrates take up different channel entrances, they pass through ABP and DBP. Sometimes, it bypasses one of the binding pockets. We noticed two DDM molecules are bound in our structure, one near the CH1 entrance (between TM8 and TM6) (**Figure 6A**) and the other binding site at the CH4 entrance (groove between TM1 and TM2) (**Figure 6B**). In an earlier study, it was shown that fusidic acid binds to the deep transmembrane binding pocket (TMD-BP) and then allosterically regulates the binding of substrates in CH1 and CH4 binding pockets. During this process, movement in TM11/TM12 is observed in the binding state protomer on substrate binding, creating a ∼1700 Å^3^ cavity to accommodate FUA, a similar cavity was created in our structure locally (Tam *et al*, 2021b).

In our structure, DDM is bound to all the binding pockets in TMD except the TMD_BP (surrounded by TM2, TM4, TM10-12). It is bound at the same positions as of DDM and fusidic acid in the TFI-LIG1 state (6zod) of the allosteric transport mechanism (**Figure 6A & B**). The proposed transport mechanism of DDM considers only the CH1 or CH4 channels at a time (Tam et al, 2021a; Eicher et al, 2012; Sennhauser et al, 2006; Tam et al, 2020; Ababou & Koronakis, 2016). In our finding we observe, the DDM is bound simultaneously to all the sites, similar to the allosteric binding of FUA in the transmembrane domain (TMD). We speculate that, upon saturating the TM binding domains with DDM, the conformational transition happens in the TMD and will extrude the substrate to the periplasmic domain (PD) by squeezing/shrinking the TMD. Similar to proton translocation or allosteric binding in TMD-BP, the conformational change has been induced from the TMD to the PD. As a result, the TMD experiences extrusion on saturation of the binding pocket while the PD is still in the binding state. In addition, the binding of multiple DDM at different binding pockets made us believe that the structure is dynamic in the transmembrane region because of the DDM binding and extrusion process. We hypothesise that the solved structural state is the post-extrusion state of MdtF, where the central pore is closed post-transportation of the DDM molecule.

Structural analysis of MdtB revealed a periplasmic domain conformation distinct from any previously reported structural states of RND transporters. Importantly, MdtB features a closed access binding pocket (ABP) and an expanded deep binding pocket (DBP) (**Figure 5A**). The transmembrane helix TM8 adopts a folded α-helical configuration resembling the extrusion state, effectively sealing the vestibule entry between TM8 and the PC2 domain (**Figure 5**). A key structural element, the G-loop-critical for substrate binding and selection, appears shorter and more rigid in MdtB compared to other RND pumps (**Supplemental Figure 7**). This rigidity is because of the reduced number of glycine residues: MdtB contains only two glycines within its G-loop, whereas typical RND transporters possess four, and MdtF contains five, enhancing its flexibility. Furthermore, MdtB lacks the conserved GFGF motif and does not have a phenylalanine residue within the loop-specifically, the highly conserved F615, known to be essential for substrate transport in RND pumps. Instead, the G-loop of MdtB is enriched in polar aliphatic residues and is structurally displaced upward, in contrast to other RND pumps where the G-loop acts as a dynamic valve between the ABP and DBP. These observations suggest that the G-loop of MdtB may play a limited role in substrate recognition and selection. To further explore this unusual feature, we examined the G-loop of MdtC, the bicomponent partner of MdtB. The MdtC loop is four residues shorter and similarly lacks aromatic amino acids (**Supplemental Figure 1**). These structural characteristics, shared by both MdtB and MdtC, suggest that the bicomponent RND system possesses an unconventional G-loop. This may lead to altered substrate specificity or facilitate distinct transport mechanisms depending on whether the pump operates in a homotrimeric or heterotrimeric state.

The asymmetric structures of RND transporters have provided critical insights into the functional rotational mechanism underlying substrate efflux, as demonstrated in AcrB bound to substrates such as doxycycline, fusidic acid, macrolides, erythromycin, and levofloxacin (Oswald et al., 2016; Ababou & Koronakis, 2016; Yu et al., 2005; Seeger et al., 2006). In addition to the canonical access, binding, and extrusion states, numerous alternative conformational states have been resolved that do not align with this classical model (Morgan et al., 2021; Kato et al., 2023; Tam et al., 2021a). Cryo-EM technique is used to captured these non-canonical conformations, including resting and intermediate states of HAE-RND transporters (Morgan et al., 2021; Zhang et al., 2021). The resting conformation, in particular, is described as an inactive state prior to channel opening and substrate entry (pre-access state). Furthermore, several intermediate states display a combination of structural features from the three traditional states, which have also been reported; one such example is BpeB and BpeF (Kato et al., 2023). These findings highlight the continuous structural rearrangements that occur during substrate translocation, making it inherently difficult to capture the full spectrum of dynamic states experimentally. Although various drug-bound structures have been resolved, the complete sequence of conformational transitions remains poorly understood. Molecular dynamics simulations of AcrB results have supported this idea, highlighting the intermediate states and demonstrating the mobility of transmembrane helices (Matsunaga et al., 2018a). These studies suggest that RND transporters are intrinsically dynamic, adopting a broad range of intermediate conformations. our study highlights the dynamic behaviour of RND pumps under defined experimental conditions by capturing transient intermediate states of MdtF and MdtB,.

In summary, this study offers significant insights into the structural and functional dynamics of RND efflux pumps, with a focus on MdtF and MdtB from *E. coli*. Contrary to previous assumptions, we showed that the homotrimeric form of MdtB is functionally active and capable of substrate transport. Importantly, we resolved previously undefined conformational states of both MdtF and MdtB, revealing novel structural features. Despite MdtF sharing high sequence similarity with AcrB and belonging to the same Acr cluster within the HAE-RND family, its structure is different from that of AcrB. In contrast, MdtB, which belongs to a different RND cluster, shows a high structural resemblance to AcrB. MdtF displays notable flexibility within its transmembrane domain, whereas MdtB features a more rigid and stable transmembrane architecture, similar to other well-characterized RND pumps such as AcrB, MtrD, AdeB, and AdeJ. Our comparative analysis highlights that these structurally divergent RND transporters employ distinct mechanisms for substrate recognition and transport.

## Materials and Methods

### Strains and constructs

The MdtF gene is PCR amplified from the *E. coli* genomic DNA, and it is cloned as pET21a (+)-vector between the NdeI and XhoI restriction sides with the C terminal octa-His tag. MdtF was expressed in the strain of *E. coli* BL21(DE3) ΔABEF cells. Kanamycin (50 μg/ml) and ampicillin (100 μg/ml) are used as a selection marker for ΔABEF cells and MdtF, respectively.

#### Multiple sequence alignment

Multiple sequence alignments were performed within the RND pumps family using the Unipro UGENE and clustal omega. Among that, based on its consensus, residues are highlighted in gradient. Besides, in comparison with the reference RND pump AcrB, residues were highlighted according to their conservation.

### Drug resistance agar plate assay

An agar plate assay was performed using an Erythromycin substrate for MdtF and MdtB. The MdtF gene is cloned into the pBAD vector for the assay, and the MdtB gene is carried in the pET21a vector. Overnight grown cells were diluted to OD600 of 1, and further serial dilution of 10-fold was prepared. The diluted sample was spotted on the Luria-Bertain (LB) agar plates. MdtF samples and pBAD vector samples are spotted on the agar plates containing 0.02% (w/v) L-arabinose, Kanamycing (50mg/ml), and Ampicillin (100mg/ml). MdtB and pET21a samples are spotted on the plates with the same composition except for arabinose. The survival of the cells was checked after 24h of incubation at 37°C.

### Protein purification

To express the wild-type efflux pumps of MdtB and MdtF, *E. coli* BL21(DE3) ΔABEF cells carrying appropriate plasmid were grown in Luria–Bertani (LB) broth. When the optical density reached 0.6, cells were induced with 1 mM IPTG and grew at 20°C for 18h. The harvested cells were pelleted by centrifugation and resuspended with resuspension buffer, 50 mM Tris pH 8.0, 500 mM NaCl, 500 µM EDTA, 0.2 mg/ml (w/v) lysozyme, and 1 mM PMSF. After lysing the cells by sonication, cell debris was removed by centrifugation (8,000 rpm, 4 °C, 20 min), and the supernatant was subjected to ultra-centrifugation (30,000 rpm, 4 °C, 2h) to separate the total membrane. The membrane pellet was solubilised in membrane resuspension buffer, 50 mM Tris pH 8.0, 500 mM NaCl, 5 mM imidazole, 10% (v/v) glycerol, 2% (w/v) n-dodecyl-β-D-maltoside (DDM), for overnight at 4 ⁰C. The remaining insoluble pellet fraction was separated by ultra-centrifugation (30,000 rpm, 4 °C, 30 mins). The supernatant was incubated with pre-equilibrated Ni-NTA agarose beads (Qiagen NI-NTA agarose beads) with the binding buffer, 50 mM Tris pH 8.0, 500 mM NaCl, 5 mM imidazole, 10% (v/v) glycerol, 0.03% (w/v) DDM (buffer A). The Ni-NTA beads were washed with wash buffer, 50 mM Tris pH 8.0, 500 mM NaCl, 50 mM imidazole, 10% (v/v) glycerol, 0.03% (w/v) DDM (buffer B) and The proteins were eluted with elution buffer 50 mM Tris pH 8.0, 500 mM NaCl, 300 mM imidazole, 10% (v/v) glycerol, 0.03% (w/v) DDM (buffer C). The eluted protein is concentrated with a 100KDa molecular weight filter Amicon spin column concentrator. The proteins are separated using size exclusion chromatography (SEC) column Superdex 200 10/300 GL with the SEC buffer, 25 mM HEPES pH 7.5, 100 mM NaCl, 0.03% (w/v) DDM. SEC-purified MdtB is incubated with amphipol A8-35 (Anatrace) in the mass ratio 1:5 at 4 ⁰C. The free detergents are removed using SM-2 biobeads. Further, amphipol A8-35 exchanged MdtB samples were separated by SEC purification.

### Negative staining TEM sample preparation and imaging

The Formvar carbon-coated copper grids from Tedpella (EM grid, 300 mesh; TedPella) were prepared by glow discharging for 30s at 20mA using the GloQube glow discharge system, Quotum. The peak fractions of MdtB and MdtF were diluted 40x times with the SEC buffer to visualise under room temperature TEM. Four microliters of diluted samples (0.1mg/ml) were placed on the glow-discharged grids and, after 60 seconds, blotted with a Whatman filter paper. Stained the sample containing grids with 1% uranyl acetate (98% uranyl acetate; ACS Reagent, Polysciences Inc. Warrington, PA, USA) followed by blotting and air drying for 2 minutes. The negatively stained samples were visualised using a Tecnai T12 electron microscope equipped with a LaB_6_ filament operated at 120 kV. At a moderately stained area, a total of 30 images were collected at a magnification of x 220,000 calibrated with 2.54 Å per pixel. The images were collected using a side-mounted Olympus VELITA (2000 x 2000) charge-coupled device camera.

### Negative staining TEM data processing

Data processing for the collected negative staining micrographs were performed with EMAN 2.1 software. In total 3000 particles were picked automatically with the EMAN software package(Tang *et al*, 2007). The particles were extracted using e2boxer.py in the EMAN 2.1 software package. The particle stack was generated with the generate stack tool in EMAN, followed by calculating Reference-free 2D class averages for the generated particle stack using e2refine2d.py. For further processing, a high pass filter of 0.2 was applied during class averaging. The particle stack was exported to Relion 2.1 to perform 2D class averaging. In the first round of 2D classification, the particles were split into 50 classes. The classes with good signal-to-noise ratios were chosen for the next round of class averaging by splitting into 30 classes. Finally, 2000 particles contributing to the best class averages were selected to execute the last round of 2D class averaging using simple_prime2D of SIMPLE 2.1 software (Reboul *et al*, 2018) with a mask diameter of 100 pixels at 2.54 Å per pixel value.

### Cryo-EM sample preparation

QUANTIFOIL 300 mesh Cu ultrathin 2 nm grids with R 1.2/1.3 size were glow discharged for 70 s at 20 mA using Quorum GlowCube. Three microliters of purified MdtB and MdtF were placed on the glow-discharged grids, followed by incubating for 10 s before blotting for 8 seconds at 100% humidity. The sample containing grids were flash frozen by quickly plunging into the liquid ethane using a FEI Vitrobot IV plunger (Thermo Fisher Scientific).

### Cryo-EM data acquisition of MdtB and MdtF using 200-kV Talos Arctica

Talos Arctica 200kV cryo-TEM (Thermo Fisher Scientific) is used for cryo-EM data acquisition of MdtB and MdtF to resolve the structure. The instrument is equipped with K2 direct electron detector (Gatan Inc.). In total of 3750 movie files were collected for MdtB and 4000 for MdtF using Latitude-S automated data acquisition tool (Gatan Inc.) (Kumar *et al*, 2021). Each movie file contains 20 frames with a total electron dose of 50 e-/Å² in the −0.75 to −2.25m defocus range. The data was collected at 45,000X magnification with the calibrated pixel size of 1.17 Å.

### Cryo-EM data processing of MdtB

The micrographs are manually screened in cisTEM software to discard the micrographs with poor signal-to-noise ratio. The best micrographs after screening using cisTEM are used for the data processing (Grant *et al*, 2018). Data was processed using RELION 3.1.2 software (Scheres, 2012). Beam-induced motion correction was carried out on the movie files using the MotionCor2 package (Zheng *et al*, 2017). The contrast transfer function (CTF) estimation of the micrographs is performed using CTFFIND 4.1.13 (Rohou & Grigorieff, 2015). For generating a 2D class average template for automated particle picking, around 10000 particles were initially manually picked, and reference-free 2D classification was performed using RELION 3.1.2 (Scheres, 2012) to choose the best 2D class averages. In Relion, 5,146,611 particles are auto picked using the reference 2D classes. Automatically picked particles were extracted with a box size of 300 pixels. Multiple rounds of 2D classification were run with the extracted particles to clean the particle stack. Around 8500 particles contributing to different orientations of MdtB were selected to generate an ab initio initial model. The best 5,42,913 particles from the dataset were 3D classified into 6 classes using the ab initio map as a reference. The 3D classification was performed with C1 and C3 symmetry separately. In asymmetric 3D classification, a 3D class with 258549 particles was observed as a well-resolved map. 3D auto refinement was carried out for the 3D map using a soft mask in RELION 3.1.2. Followed by performing per-particle defocus refinement with beam tilt correction, anisotropic magnification correction and per micrograph astigmatism fitting. Finally, the corrected particles were polished in Relion using Bayesian polish. The polished particles were subjected to final 3D refinement and sharpening. The 3D classification with C3 symmetry generated a class with a well-resolved map contributed by 327240 particles. Refinements and post-processing are carried out similarly to the C1 symmetry map. The structural differences between symmetric and asymmetric reconstruction were analysed after superimposing both the cryoEM maps in UCSF Chimera and UCSF ChimeraX (Pettersen *et al*, 2004; Goddard *et al*, 2018).

### Cryo-EM data acquisition of MdtF using 300-kV Titan Krios

Cryo-EM data were acquired using a 300-kV operated Titan Krios (Thermo Scientific) microscope equipped with a Falcon direct electron detector (Thermo Fisher Scientific). Micrographs were automatically collected using EPU software (Thermo Scientific) at a nominal magnification of 75,000x with an effective pixel size of 1.07 Å at specimen level. These recorded movies have 30 frames, with a total exposure dose of 30.39 e^-^ /Å^2^ at a defocus range of −1.5 to 2.7 μm. A total of 3000 micrographs were collected for further data processing.

### Cryo-EM data processing of MdtF

The entire single particle analysis (SPA) data processing was performed in RELION 2.1 (Scheres, 2012) and cryoSPARC (Punjani *et al*, 2017). After screening the micrographs using cisTEM, micrographs with poor signal-to-noise ratio were discarded (Grant *et al*, 2018). The beam-induced motion correction was carried out on the movie files using MotionCorr2 software (Zheng *et al*, 2017). The contrast transfer function (CTF) of micrographs was estimated using CTFFIND 4.1.13 (Rohou & Grigorieff, 2015). Initially, a blob picker was used to pick particles from the subset of micrographs to generate a template for picking. 1,00,000 particles were extracted with a 300-pixel box size and were sorted by multiple rounds of 2D classification. The final 2D class average was used as a template to pick particles from the full data set. In total, 1815916 particles were picked. Multiple rounds of 2D class averaging were performed to select 1025190 particles, which had a good signal-to-noise ratio. The good particle set was transferred to cryoSPARC for further data processing (Punjani *et al*, 2017). Ab initio reconstruction was performed with C3 symmetry, and the map was used as a reference with a low-pass filter of 20 Å for further non-uniform refinement. After non-uniform refinement with a mask, the final map was sharpened with a −150 B-factor. The final resolution of the refined MdtF map with C3 is determined to be 2.8 Å. The good particle set of 752103 is used for Ab initio 3D reconstruction with C1 symmetry and split into three. The maps are further refined by performing non-uniform refinement with the 20 Å low pass filtered initial models. The final maps are sharpened with a B-factor of −100. The overall pipeline of EM data processing is shown in the **supplemental figure 3**.

### Model building and analysis

MdtB and MdtF models are generated using alpha fold web tool. The generated models are fitted in the map using ChimeraX tool (Goddard *et al*, 2018). FlexEM tool from CCPEM was used to fit the flexible secondary structural regions. The models are real space refined using the Phenix refinement program and corrected using Coot (Adams *et al*, 2010). The models are validated with Phenix and Coot software for map-to-model validation, Ramachandran outliers, and clashes. Substrate transport channels and pores were analysed for MdtF and MdtB using MOLEonline webtool and CASTp webtool.

## Acknowledgements

We thank the Advanced Center for Cryo-Electron Microscopy Facility, IISc, Bangalore, for data collection in Talos Arctica 200kV. We acknowledge the Department of Biotechnology, Department of Science and Technology (DST) and Ministry of Human Resource Development, India for funding the cryo-EM facility at IISc-Bangalore. We acknowledge the DBT-BUILDER Program (BT/INF/22/SP22844/2017) and DST-FIST (SR/FST/LSII-039/2015) for the National Cryo-EM facility at IISc, Bangalore. We thank Dr. Vinothkumar KR and Dr. Sucharita Bose for data collection at the National Electron Cryo-Microscopy Facility, NCBS. Research in the manuscript was supported by STARS (SERB-STR/2022/000006) and SERB CRG/ 2022/002674 fellowships. RB is funded by DBT-Wellcome Trust early career fellow (IA/E/19/1/504978). SP is a graduate student funded by the Council of Scientific and Industrial Research (CSIR) fellowship. CFR is a graduate student funded by the Indian Institute of Science (MHRD) fellowship. We thank Dr. Saikat Chowdhury for the critical scientific discussion.

## Author contributions

Somnath Dutta: Project design and administration, resource support, supervision, cryo-EM data collection and processing, analysis, manuscript writing, reviewing and editing.

Surekha Padmanaban: Sample preparation, biochemical assay, NS-TEM and CryoEM data processing, model building, data analysis, manuscript writing, review, and editing.

Clayton Fernando Rencilin: Sample preparation, NS-TEM and CryoEM data collection and processing, manuscript reviewing and editing.

Rupam Biswas: Project design, sample preparation, initial cryoEM data processing, manuscript review.

## Deposited Data Accessibility

Cryo-EM density maps of the asymmetric structure of MdtF and symmetric structure of MdtB were deposited in the Electron Microscopy Data Bank (EMDB), EMD-63315, EMD-63321. The PDB structure of MdtF ID is 9LYJ, and MdtB ID is 9LYQ.

