## Supplemental file for "Cryo-EM reveals the structural heterogeneity and conformational flexibility of multidrug efflux pumps MdtB and MdtF"

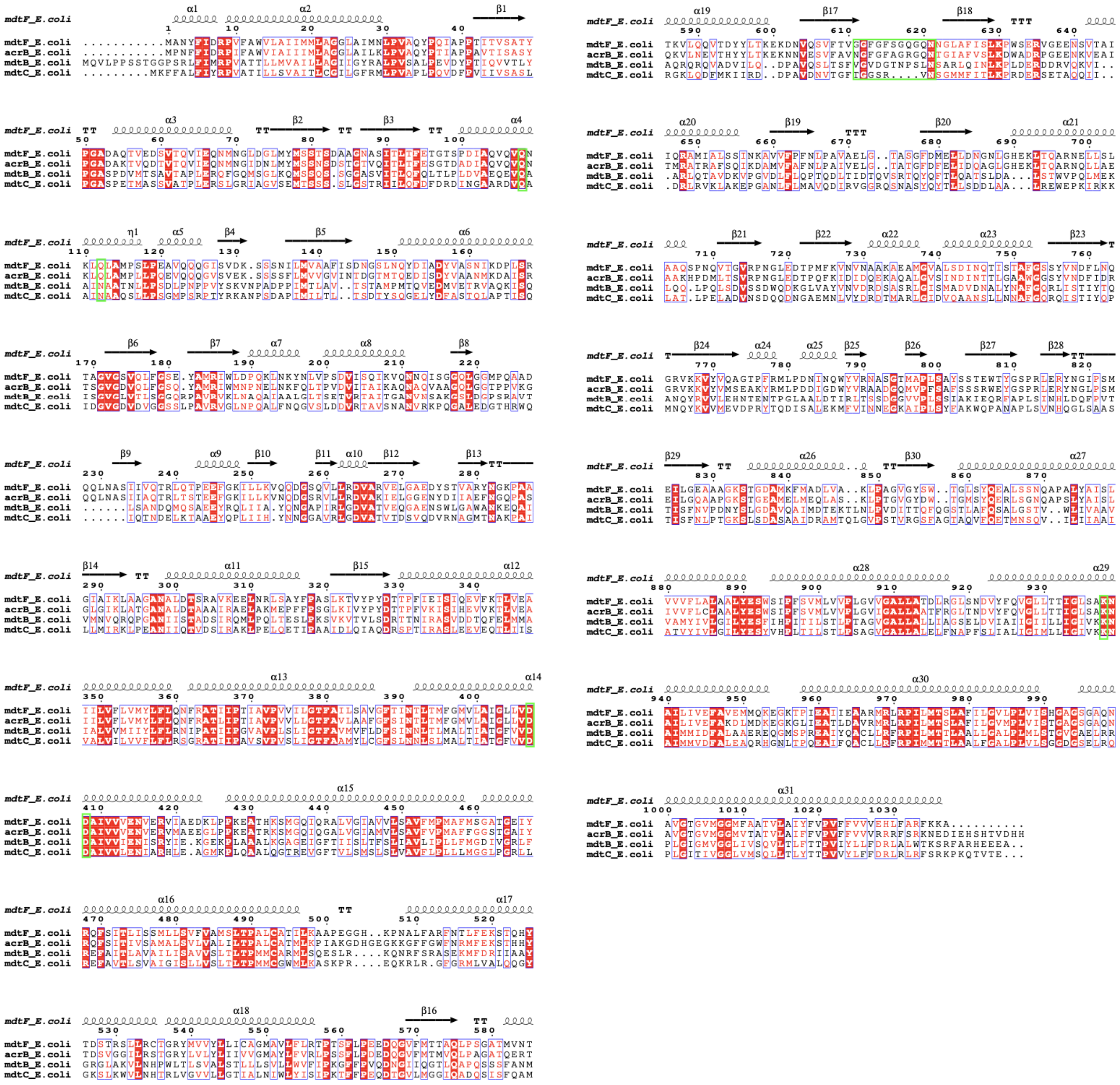

**Supplemental Figure 1:** Multiple sequence alignment of RND pumps.

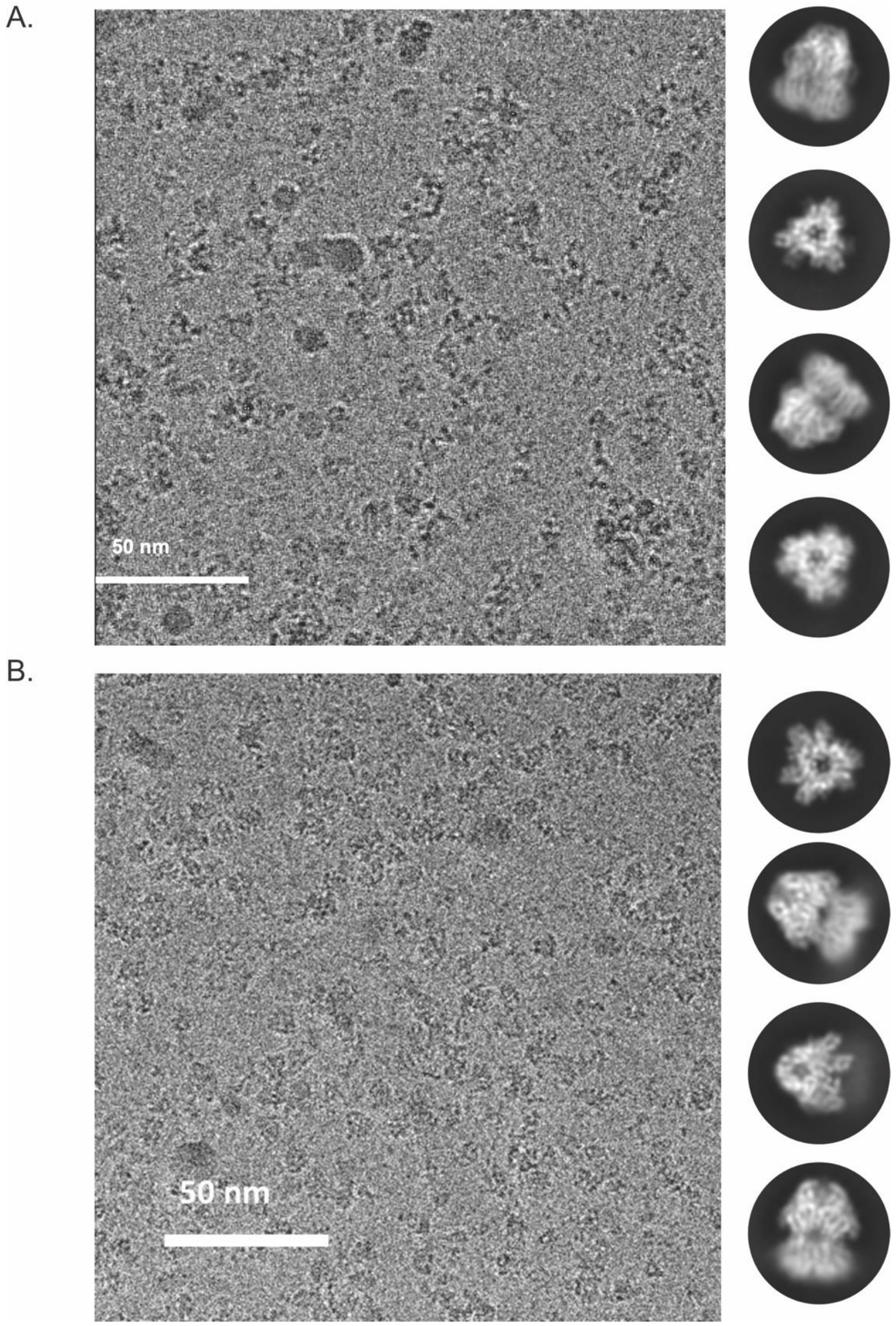


**Supplemental Figure 2:** A, Cryo-EM micrograph and 2D class average of MdtB. B, CryoEM micrograph and 2D class average of MdtF.

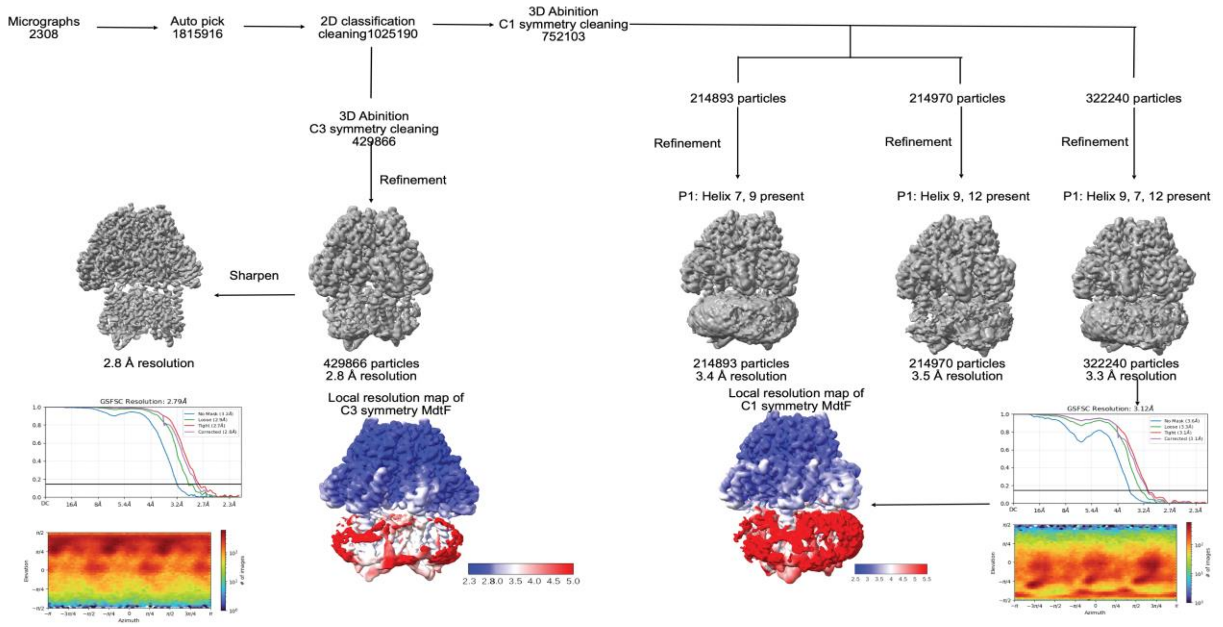


**Supplemental Figure 3:** Cryo-EM data processing workflow of MdtF

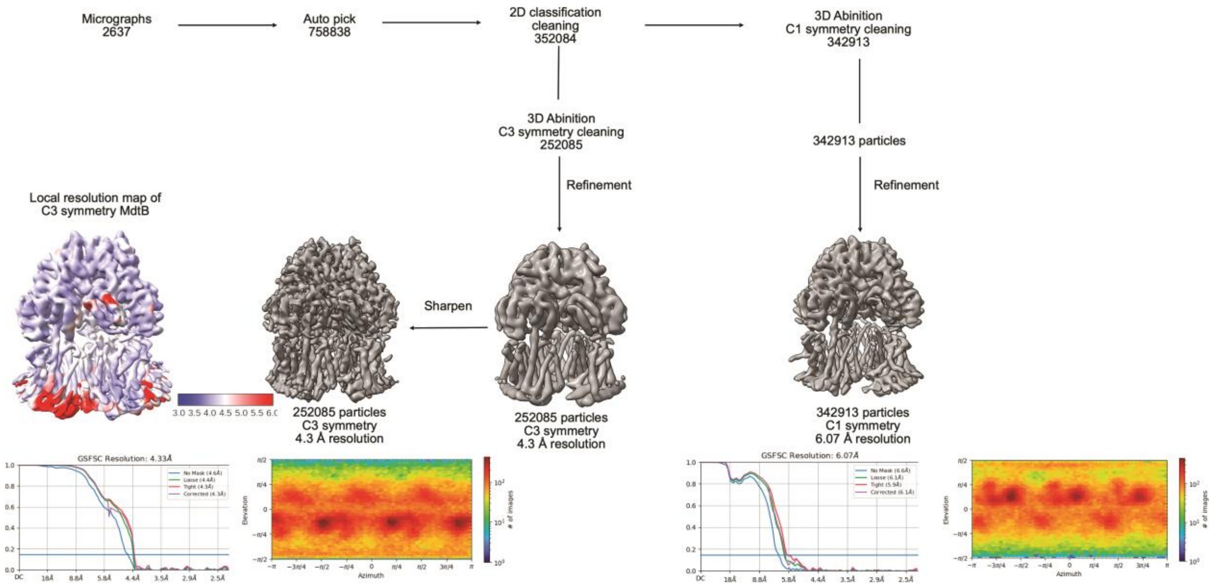


**Supplemental Figure 4:** Cryo-EM data processing workflow of MdtB

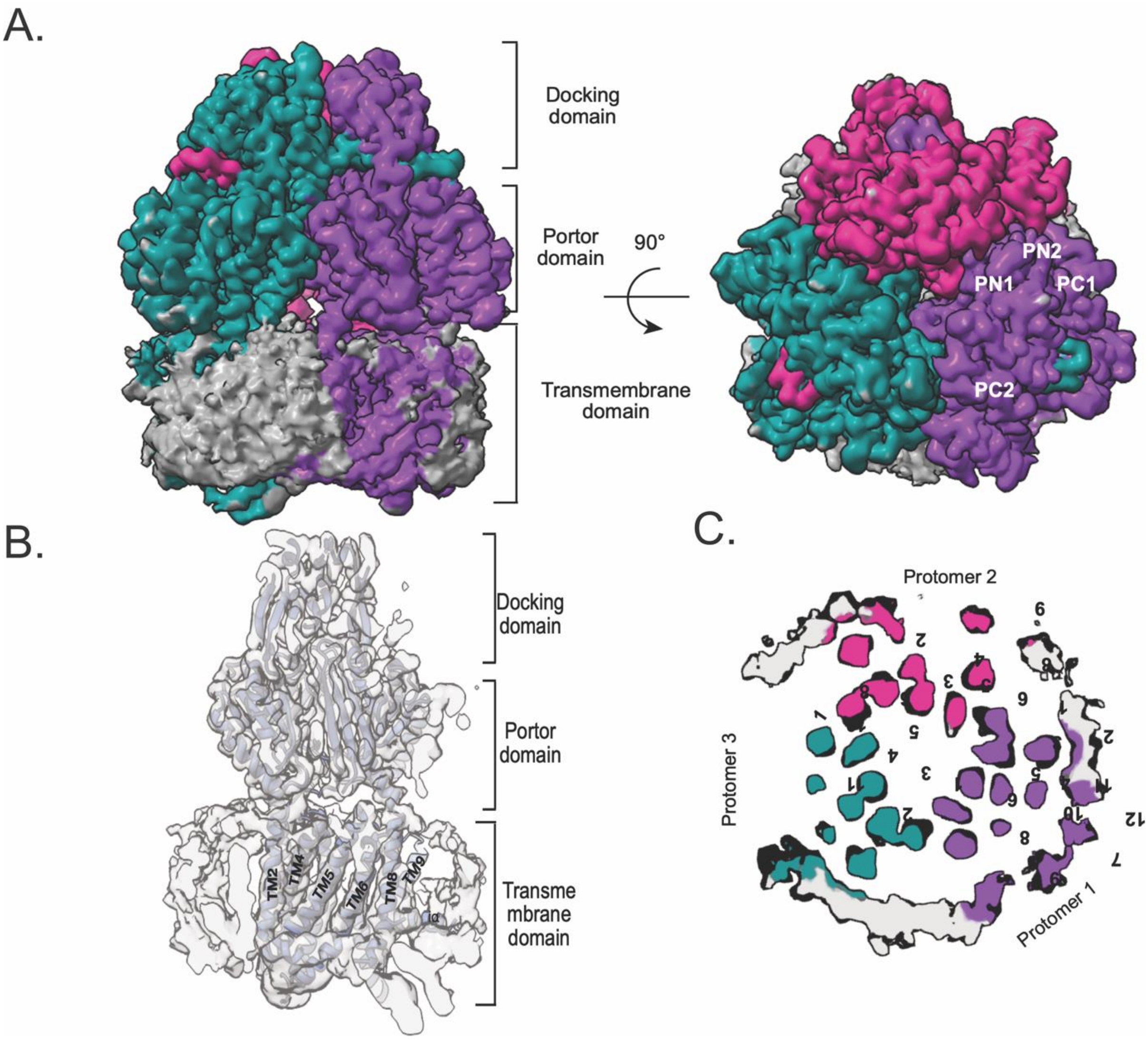


**Supplemental Figure 5:** Asymmetric reconstruction of MdtF. A, Protomer 1 with all the resolved transmembrane helixes solved the structure of the peripheral helixes in MdtF. B, Top view of the resolved helix of protomer 1. C, MdtF model fit in the map.

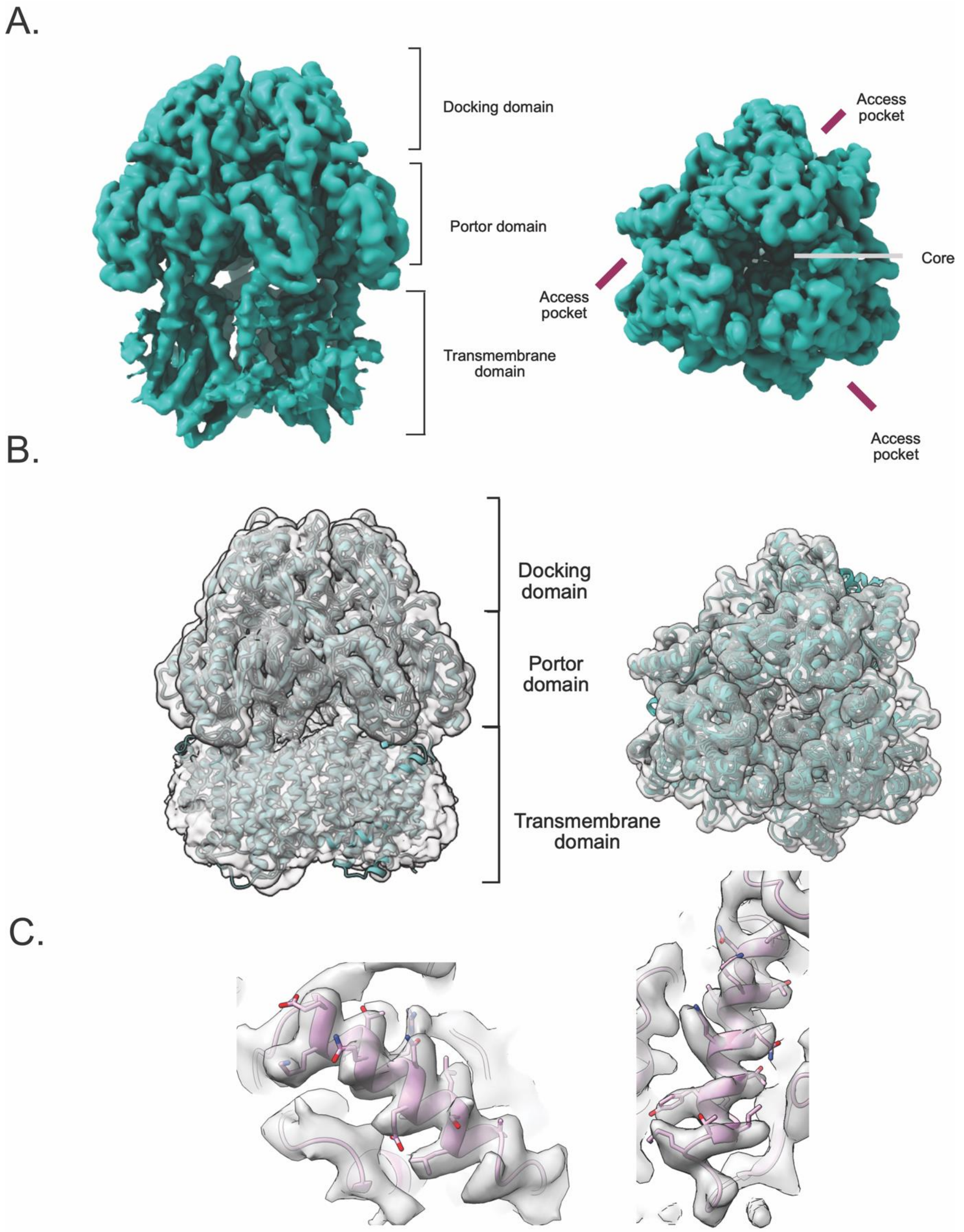


**Supplemental Figure 6:** MdtF structure solved with 200kev Talos Arctica: A, MdtF cryoEM map resolved at 3.3 Å resolution. B, MdtF model fitting in cryoEM map. C, Sidechain fitting of the helix.

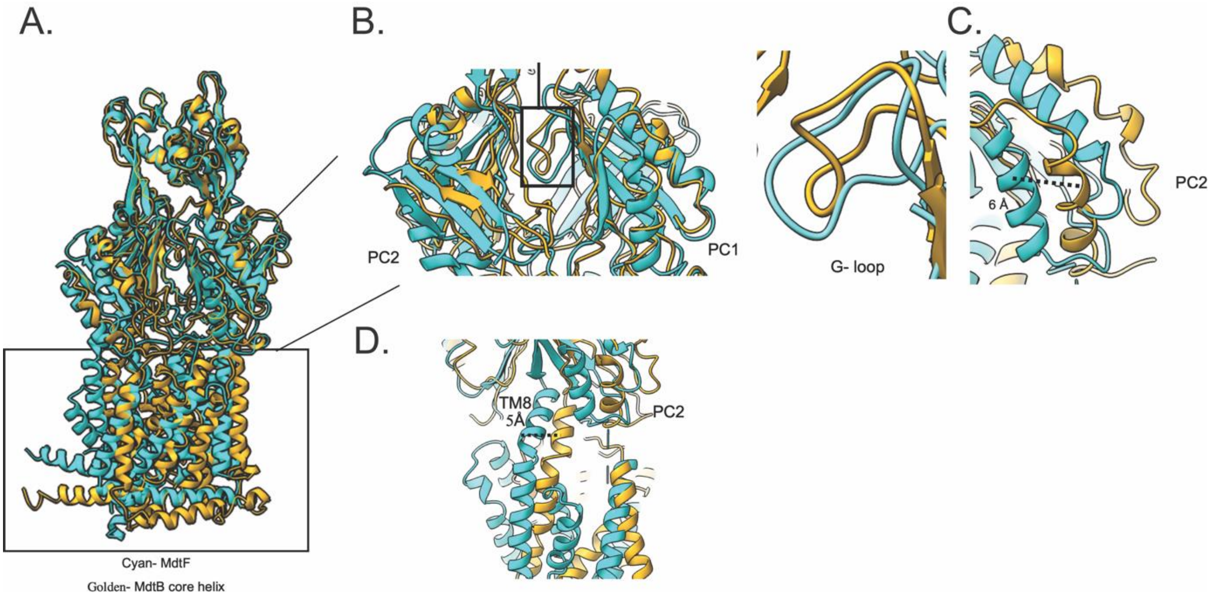


**Supplemental Figure 7:** Comparison of MdtF with MdtB. A. MdtF superimposed on MdtB. B, G loop of MdtF and MdtB. C. PC1 movement comparison between both the proteins. D. TM8 and PC2 comparison between MdtF and MdtB.

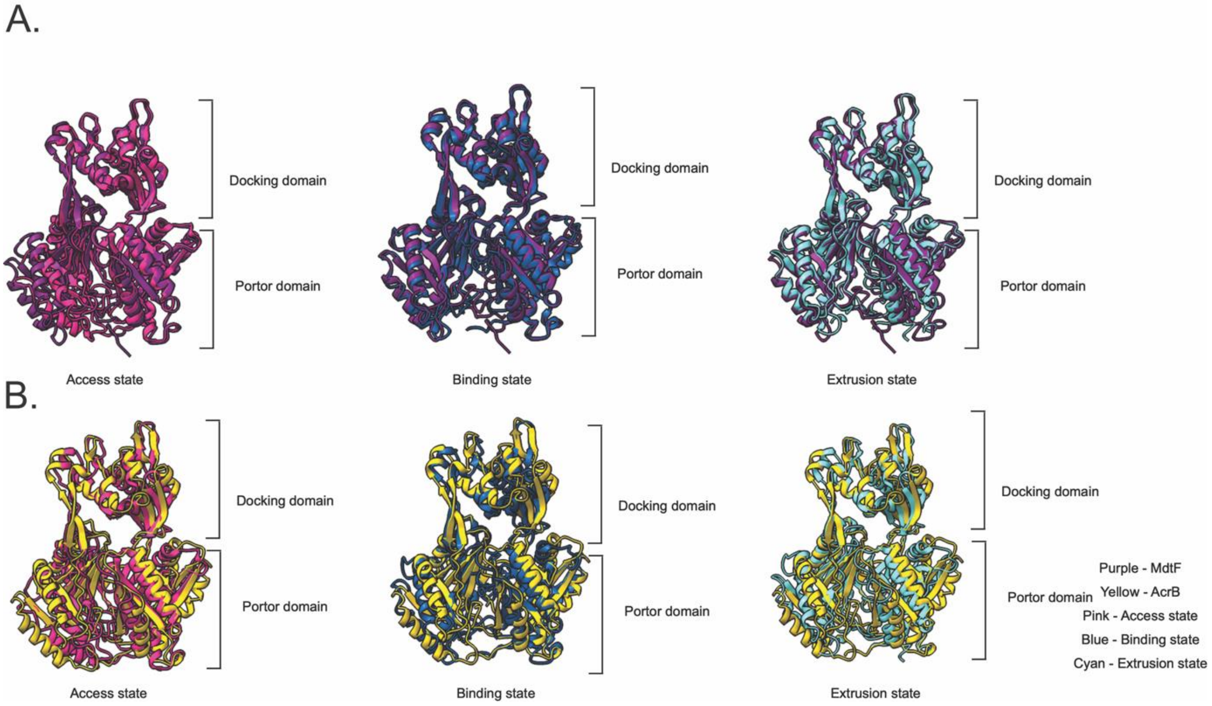


**Supplemental Figure 8:** Comparison of MdtF and MdtB with different states of RND pump. A, Comparison of access state, binding state, and extrusion state of MdtF. B, Comparison of access state, binding state, and extrusion state of MdtB.

**Table S1**: **Cryo-EM Data collection, image processing and refinement for MdtF**

| **Data Collection and Processing** | **MdtF-300kV** (asymmetric) |
| --- | --- |
| Magnification | 75,000x |
| Voltage | 300kV |
| Electron exposure ($e^-$/Å$^2$) | 30.39 $e^-$ /Å$^2$ |
| Defocus range (µm) | -1.5 to 2.7 µm |
| Pixel size (Å) | 1.07 Å |
| Symmetry Imposed | C1 |
| Number of Particles | 322240 |
| Map Resolution (Å) | 3.3Å |
| FSC threshold | 0.143 |
| Map Resolution Range | 2.5-5.0Å |
| Map Sharpening B-Factor (Å$^2$) | -150 |
| Number of Movies | 3000 |

**Table S2: Cryo-EM Data collection, image processing and refinement for MdtB**

| **Data Collection and Processing** | **MdtB – 200kV** |
|---|---|
| Magnification | 42900x |
| Voltage | 200kV |
| Electron exposure ($e^-/Å^2$) | 50 e-/Å² |
| Defocus range (µm) | -0.75 to -2.25µm |
| Pixel size (Å) | 1.17 Å |
| Symmetry Imposed | C3 |
| Number of Particles | 252085 |
| Map Resolution (Å) | 4.3 Å |
| FSC threshold | 0.143 |
| Map Resolution Range | 3.0-5.0 Å |
| Map Sharpening B Factor ($Å^2$) | -150 Å |
| Number of Movies | 3750 |
